# Spectrally Resolved Single Molecule Orientation Imaging Reveals Direct Correspondence between Polarity and Order Experienced by Nile Red in Supported Lipid Bilayer Membrane

**DOI:** 10.1101/2024.06.21.600028

**Authors:** Aranyak Sarkar, Jyotsna Bhatt Mitra, Veerendra K. Sharma, Vinu Namboodiri, Manoj Kumbhakar

## Abstract

Molecular level interaction among lipids, cholesterol and water dictates nanoscale membrane organization of lipid bilayers into liquid ordered (Lo) and liquid disordered (Ld) phases, characterized by different polarity and order. Generally, solvatochromic dyes easily discriminate polarity difference between Lo and Ld phases, whereas molecular flippers and rotors show distinct photophysics depending on membrane order. In spite of progress in single molecule spectral imaging and single molecule orientation mapping, still direct experimental proof linking polarity with order sensed by the same probe eludes us. Here, we demonstrate spectrally resolved single molecule orientation localization microscopy to connect nanoscopic localization of probe on bilayer membrane with its emission spectra, three-dimensional dipole orientation and rotational constraint offered by the local microenvironment and highlights the beautiful correspondence between polarity and order. This technique has the potential to addres nanoscale heterogeneity and dynamics, especially in biology as well as material sciences.

**TOC GRAPHICS:** 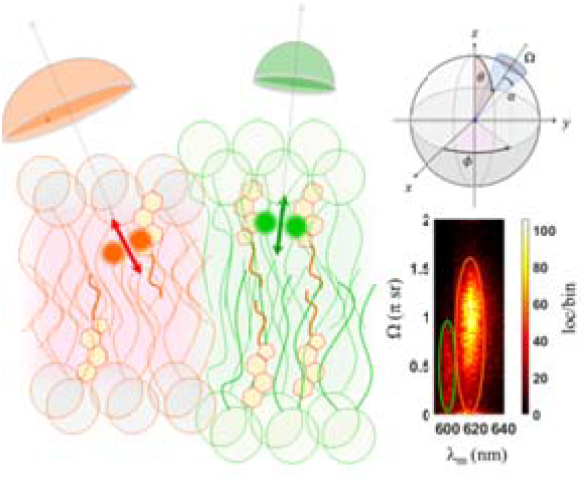

## TEXT

Membranes are dynamic interfaces made up of lipid self-assemblies in bilayer architecture by partitioning water molecules away from the hydrophobic lipid tails.^1-3^ This special structure allows compartmentalization of cells and cell components, yet fluid enough to allow lateral diffusion of constituents and tethered membrane proteins and cargos.^4, 5^ Molecular level interactions among component lipids, cholesterol and proteins in warding off extracellular water matrix define submicron membrane organization and its distinct bio-physicochemical characteristic.^6-8^ Hence, precise understanding of membrane structure and organization in relation to its intrinsically heterogeneous molecular interactions and thus environment across the length and depth of the two dimensional layer in assessing its unique behavior have captivated our attention.

Generally, model-membranes as a gross substitute to cell membrane are extensively explored to unravel the preferential molecular interaction of lipids and the origin of inhomogeneous lipid distribution in cell membranes. Investigations of model-membrane organization reveal cholesterol (and sphingomyelin) dependent lateral segregation of membrane in liquid ordered (Lo) phase formed by saturated lipids and cholesterol, surrounded by liquid disordered (Ld) phase made up of mostly unsaturated lipids (see **Schematic 1a**).^9^ Here, the planarity of rigid sterol rings of cholesterol prefers saturated lipids with linear hydrocarbon chains over twisted chains of unsaturated lipids, resulting in extended conformation for saturated hydrocarbon chains and ordering.^9, 10^ Such selectivity of cholesterol for the Lo phase is also explained considering cholesterol hydrophobicity through umbrella effect.^11^ Strongly hydrated polar phospholipid head groups of constituent lipids, e.g. phosphatidylcholine (PC) in 1,2-dioleoyl-sn-glycero-3-phosphocholine (DOPC) or in 1,2-dipalmitoyl-sn-glycero-3-phosphocholine (DPPC), preferentially shield the non-polar cholesterol molecules from unfavorable cholesterol-water contact, while hydrocarbon sterol portion of cholesterol is embedded within the nonpolar core of the lipid membrane.

**Scheme 1.**
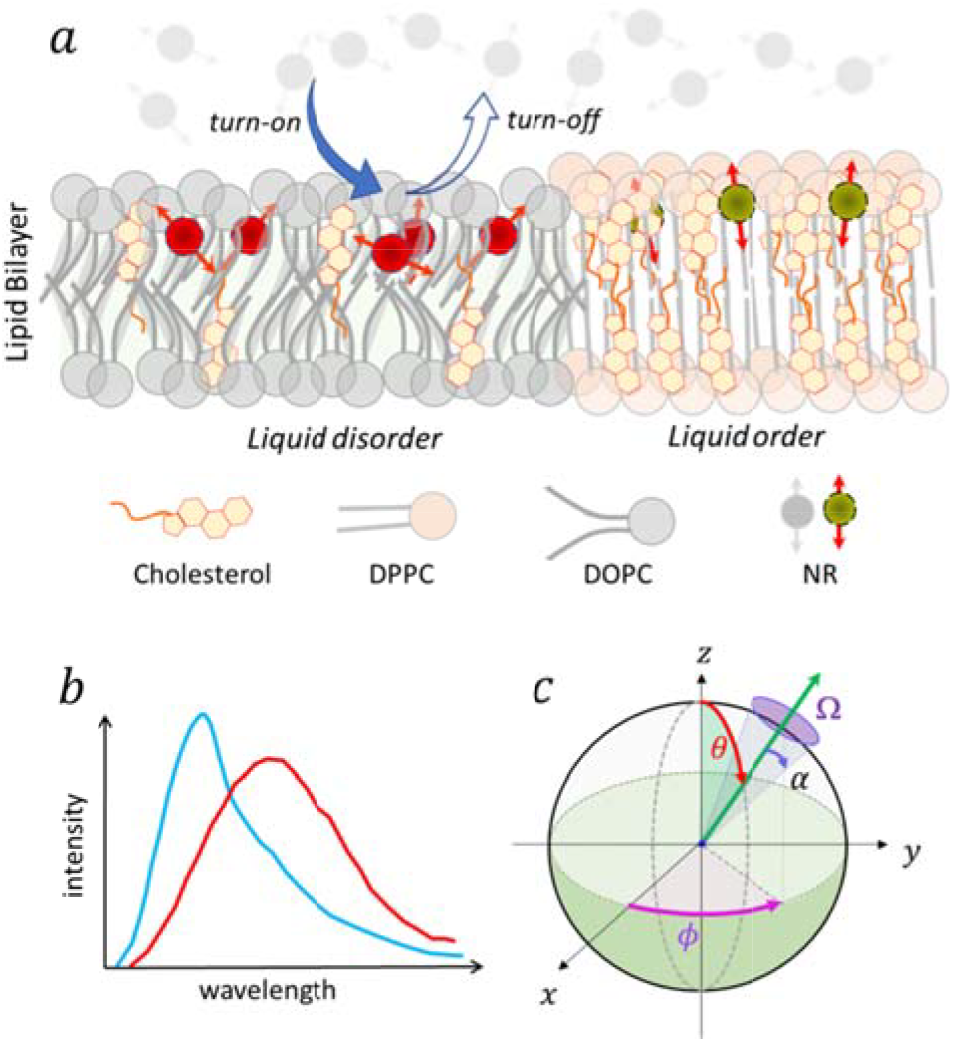
(a) Graphics depicting sensing by binding-activation of environment sensitive fluorescent probe NR on phase segregated lipid bilayer membrane of DOPC/DPPC/Cholesterol. Unsaturated lipid (DOPC) forms liquid disorder phase while saturated lipid (DPPC) forms the liquid order phase. Cholesterol predominantly resides in the order phase. (b) Cholesterol rich ordered phase is expected to show less hydration and thus blue shifted emission for environment sensitive probe NR than in th cholesterol poor disordered region. (c) Dipole orientation in laboratory frame, where z-axis is the optical axis, polar angle *θ* is the angle of the dipole (green arrow) relative to the optical axis (0 ≤θ ≤ 90°) and *ϕ*is the azimuthal angle (0 ≤*ϕ* ≤ 360°). Rotational constraint experienced by the dipole inside membrane is measured by the wobbling area (*Ω*) for the shown half-cone angle (*α*). Dipoles residing in the liquid disorder phase is expected to be relatively rotationally labile than in the liquid order phase. Therefore, local environment of a dipole inside membrane can be defined by its orientation angles (*θ,ϕ*), wobbling area (*Ω*) and spectral centroid (*λ*_m_) other than its localization (*x,y*) in the Lo or Ld phase.

Though variou techniques have been applied to explore lipid organization and its microenvironment, it is the fluorescence based methods (mainly imaging and tracking)^7, 12-21^ that drew much attention in the study of reaction field presented by these self-assembled amphiphilic lipid molecules. This is primarily due to its noninvasive nature, improved spatial resolution and easy adaption to live cell study. In fluorescence spectroscopy, broadly two class of push-pull fluorophores^22-26^ are generally employed to study lipid bilayers, namely solvatochromic (polarity sensitive) dyes^26-31^ and flipper probes (intramolecular rotors as membrane tension probe),^32-36^ where magic wand is probe’s aptitude to distinguish Lo and Ld phases by altering either its spectral position (i.e. emission color),^7, 37^ excited state lifetime,^25, 38-40^ generalized polarization (GP) parameter^41^ or spectral intensity ratio^14^ linked to lipid membrane organization. Solvatochromic probes display altered emission maxima in response to changes in its local microenvironment, that is degree of membrane hydration and the magnitude of solvation dynamics of water at the solubilization site.^2^ These polarity sensitive push-pull dyes undergo intramolecular charge transfer in the excited state from the electron donor moiety to the electron acceptor moiety.^42^ Greater stabilization of these excited dipole states in polar solvents results shift in emission maxima towards longer wavelengths (see **Schematic 1b**). In general, Lo phase, where cholesterol preferentially gets solubilized, is less polar (i.e. less hydrated) by excluding polar water molecules and relatively more ordered than Ld phase.^43^ The GP value of probes like Laurdan, Pro12A, NR12S, NR12A, etc. is often linked to membrane polarity & fluidity, where low GP usually assigned to high polarity & fluidity or low cholesterol content in the membrane.^2, 36^ Besides traditional average GP value measurement, there are other elegant and advance ways to describe local order and molecular level interactions experienced by these smart-membrane probes, for example, rotational restrictions by fluorescence anisotropy,^36^ lateral diffusions by imaging FCS,^4^ single particle tracking^12^ and 3 dimensional alignment of probe (see **Schematic 1c**) inside membrane by dipole orientation imaging.^20, 21, 31, 44-47^

Above and beyond, many push-pull probes have distinct dependency of excited state lifetime on polar protic nature of surrounding medium along with configurational rigidity.^36^ Polarity dependent spectral shift in membranes corroborates well with the change in fluorescence lifetime.^14, 40^ Planarizable mechano-sensitive fluorescent flipper molecules are also very promising to image membrane tensions to distinguish Lo and Ld phases based on its significant change in fluorescence lifetime in tune with the forces exerted by the lipid membrane.^32, 35^ However arguably, these two key aspects of membrane biophysics (i.e. polarity and order) were never measured simultaneously and independently to cross check their direct connection, in spite of large number of membrane studies with fluorescent probes.

Advances in super-resolution fluorescence microscopy (SRM) has allowed spatial segregation of Lo and Ld phases of lipid bilayer membranes with nanoscale resolution using various solvatochromic probes.^48^ Progress with single molecule (SM) spectral imaging^17, 19, 49^ and specifically its implementation into SRM by many groups^16, 18, 50-56^ empowered investigations of molecular interaction & local environment inside membranes. SM super-resolution spectral imaging has clearly revealed dissimilar polarity (and thus intuitively orders) for Lo & Ld phases in live cell membranes establishing compositional heterogeneity.^7^ It is the lipid-water interaction (and more specifically few water molecules directly hydrating the lipid head group) that play the pivotal role in modulating membrane architecture and lipid diffusion behavior.^8^ Independent developments of sophisticated imaging techniques to map three dimensional orientation (*θ,ϕ*) and orientational mobility (*Ω*) experienced by SM dipoles (refer **Schematic 1c**) makes it feasible to directly quantify membrane order with SRM.^20, 21, 31, 45-47, 57^ And in fact, recently reported orientation map with SM orientation localization microscopy (SMOLM) using solvatochromic probe NR clearly distinguish differential order in Lo and Ld phases and nicely compliments earlier investigations of nanoscale compositional heterogeneity in membrane from spectral imaging.^20, 21, 31^ Therefore, certainly molecule’s 3D dipole orientation with wobbling area gives alternate proposition to explore local environment and its interaction, besides spectral characteristic of solvatochromic probes. So it will be interesting to know directly how polarity is related with membrane order in the Lo and Ld phases. However, there is no direct experimental evidence to establish link between water penetrations in bilayer-membrane (i.e. membrane polarity) with rotational constraint (i.e. lipid order) experienced by a probe. Additionally, polarity sensed by a solvatochromic probe and order sensed by flipper/rotor probe may or may not be correlated depending on their inherent photo-physics and local solubilization site in bilayer membrane architecture. Therefore, extraction of rotational constraint along with environment sensitive spectra of the same molecule with nanometer localization is imperative to answer the above fundamental question. Even with the exceptional expansions in the field of multi-dimensional super-resolution imaging,^58, 59^ the realms of dipole orientation imaging and spectral imaging are still distinctly apart. Incorporation of these two powerful techniques into a unified single nanoscopy delineates the limitless possibility of investigating not only membrane local environments but in general any microscopic heterogeneous system. And the present contribution is a logical step towards it.

To be noted, a preliminary attempt was made with quantum materials to bridge this gap by incorporating additional 2-channel X-& Y-polarized detection of emission signal along with spectral imaging for in-plane dipole angle (*ϕ*) measurement from their integrated intensity.^60^ But then again this approach is inadvertently inadequate to determine the most important parameters, i.e. the out-of-plane dipole angle (*θ*) and the intrinsic orientational mobility (i.e. wobbling area, *Ω*) as elegantly done in independent SMOLM setups using rigorous mathematical fitting of polarized SM point spread functions (PSFs).^20, 47, 57^ Expansion of the above detection scheme with 4-channel (0°, 45°, 90° and 135°) polar detection^46, 61^ instead of 2-orthogonal polar channels for extraction of all the orientation parameters (*θ, ϕ* and *Ω*) besides spectra (*λ*) is not endorsed here as it will excessively burden the limited available photon budget from SMs. In this present case, single detection channel with compact modified PSF suitable for simultaneous orientation and localization microscopy with simple optical layout is most appropriate. With this perspective, we embark on developing spectrally resolved SMOLM (SR-SMOLM) to demonstrate such possibility of nanoscopic localization of every stochastic single fluorophore binding event on membrane simultaneously with its emission spectra, 3-dimentional dipole orientation in space and rotational mobility offered by the local environment (*cf*. **Schematic 1**). Here we show the correlation between polarity and rigidity of Lo and Ld regions experienced by a common environment sensitive membrane probe **NR** (*cf*. **Schematic S1**), known to show differential emission maxima depending on solvent environment, in supported lipid bilayer (SLB) membrane of DOPC/DPPC/Cholesterol^62^ (35:35:30) (*cf*. **Schematic S2**).

Parallel detection of orientation and spectra of single molecules is preferred over sequential imaging, as the later compromises on correlation between local polarities encountered by an individual molecule with its rotational constraint. Here we demonstrate binding-activated super-resolution single molecule localization microscopy (SMLM) of NR on membrane, constructed using wide-field total internal reflection excitation based points accumulation for imaging in nanoscale topography (PAINT).^63^ In our SR-SMOLM setup, we employed two detection paths (*cf*. **Figure S1**). The emission signal collected from a single molecule is divided into two components - one for spectral imaging and another for orientation imaging. Due to the limited number of photons emitted by a single NR molecule during its binding to membrane further splitting into X-pol and Y-pol channels for SM PSF fitting^20, 21, 42^ is avoided. Additionally, conventional polarized PSFs exhibit best sensitivity for in-plane (*θ*∼90°) dipoles, but when predominantly dealing with out-plane dipole configurations (*θ* ≪ 90°, expected for NR dipoles in SLB), use of phase modulation optics^47^ is the preferred option to enhance precision of extracted orientation parameters. Simultaneous estimation of 3D position of a single dipole along with its orientation and degree of rotational constraint has been elegantly demonstrated using excitation modulated Vortexed PSF fitting for reorienting adhered intercalator dye on *λ*-DNA.^47^ In our approach, a Vortex phase plate (VPP) is positioned in the Fourier plane (FP) of the 4*f*-detection system (*cf*. **Figure S1**). The resulting Vortex PSF eliminates the need for polarization splitting and maintains a compact PSF size, facilitating easy integration with localization microscopy techniques. Employing a vectorial PSF fitting routine and calibrating for field-dependent aberrations (*cf*. **Figure S2**), we could achieve orientation and position estimation within 30% of the Cramér-Rao lower bound (CRLB) (*cf*. **Figure S3**).

For spectral imaging, a prism-based dispersion of SM PSFs^19, 53^ is implemented in the setup which is recorded by the same EMCCD camera (*cf*. **Figure S1**). To pinpoint wavelength positions on the dispersed light patch, a 10 nm narrow bandpass filter is employed, presenting a perfect 2D Gaussian PSF (*cf*. **Figure S4**). The center of this PSF signifies a specific wavelength corresponding to the bandpass used in the spectral image, and once the spectral position is identified, the entire spectrum of the molecule is computed using a pixel shift-wavelength calibration curve (see **Supporting Information**). This calibration plot is established through a series of narrow bandpass filters, as illustrated in **Figure S4**. For spectral calculation, a transformation matrix is initially constructed to map the spectrum of each individual molecule to its corresponding localization coordinates, and thus linked to its orientation (*cf*. **Schematic S3**). Consequently, this setup (as displayed in **Figure 1**) allows simultaneous recording of the spectrum and orientation parameters of SMs. The Vortexed PSFs and corresponding spectral dispersions were fitted to extract orientation (θ,ϕ,Ω) and localization (*x,y*) parameters and spectra of each SMs. In our setup with an average 3000 photons per NR molecule, we could achieve a precession of around 10 nm for 2D localization, around 4° for orientation and ∼ 2 nm spectral resolutions (see **Figure S3**). Spectral centroid (*λ*_m_) is calculated from the distribution of spectral intensity and was subsequently used to construct *λ*_m_-map, indicator of local environment polarity. Polar angle defines how well individual NR dipoles are oriented along the normal to the membrane plane, which is in a way reflects level of configurational alliance NR molecules forge with cholesterol and lipid chains, while wobbling area indicates dipole’s rotational mobility at that site. So both, polar angle and wobbling area are the articulation for the extent of lipid order reported by individual NR molecules.

**Figure 1.**
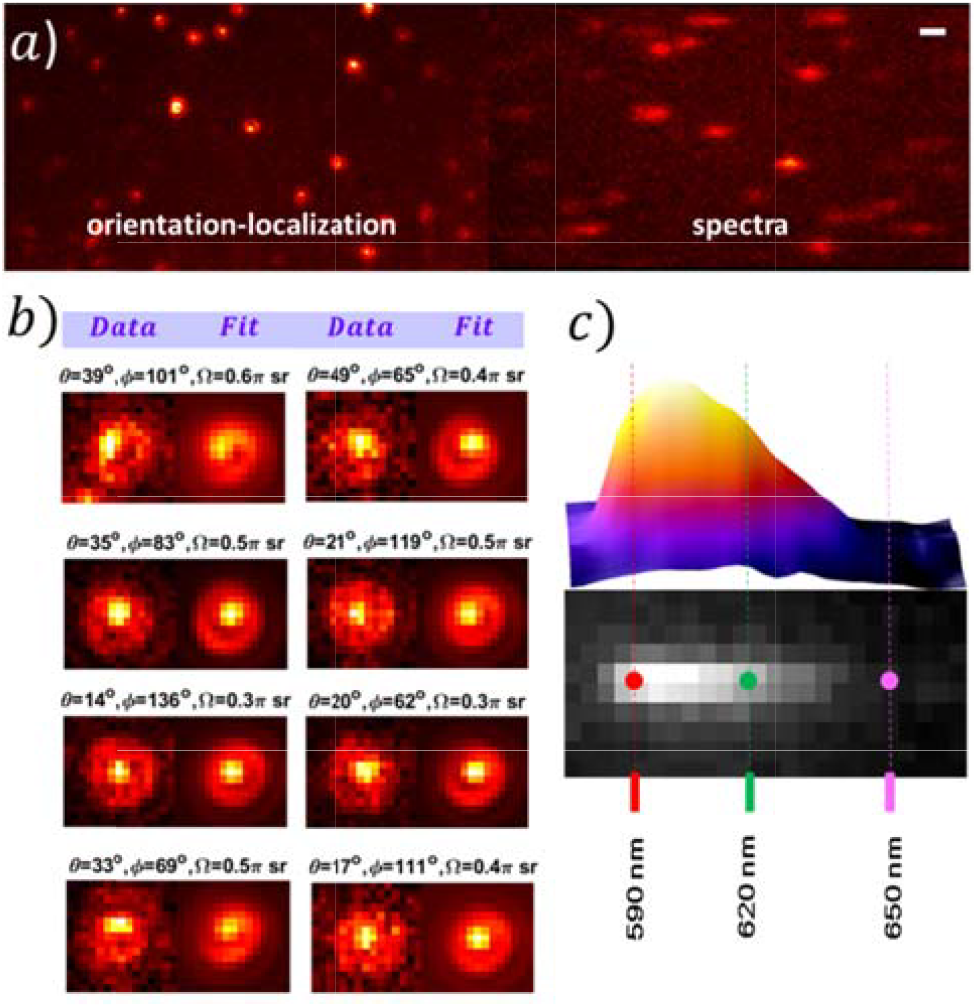
(a) A representative single image frame from stack of 45,000 frames of NR molecules on membrane showing Vortexed PSF of SMs and its spectral dispersion recorded with SR-SMOLM setup. Image recorded at frame rate of 20 Hz. (b) Raw and simulated Vortexed PSF of few selected SMs with fitted orientation parameters (*θ,ϕ,Ω*). (c) SM emission spectra with calibrated wavelengths axis. Scal bar: 1 *μ*m.

The representative super resolved image comprising SMLM, polar angle (*θ*), wobbling area (*Ω*) and emission centroid (*λ*_m_) map of SLB obtained with NR are shown in **Figure 2**. As reported in literature, unlike MC540 (see **Supporting Information**) SMLM image for NR (shown in **Figure 2a-d & 3a-d**) does not show noticeable high contrast between Lo and Ld phase in terms of localization density.^64, 65^ But, appreciable enough to segregate low and high localization regions in tandem with *θ* –map, *Ω* -map and *λ*_m_ –map. Distribution of spectral centroid constructed from all the detected SMs (around 300,000 in **Figure 2**) clearly show two distinct sub-populations around 600 nm and 620 nm, though corresponding plots for polar angle and wobbling area shows very broad distribution (*cf*. **Figure 2e-g**).

**Figure 2.**
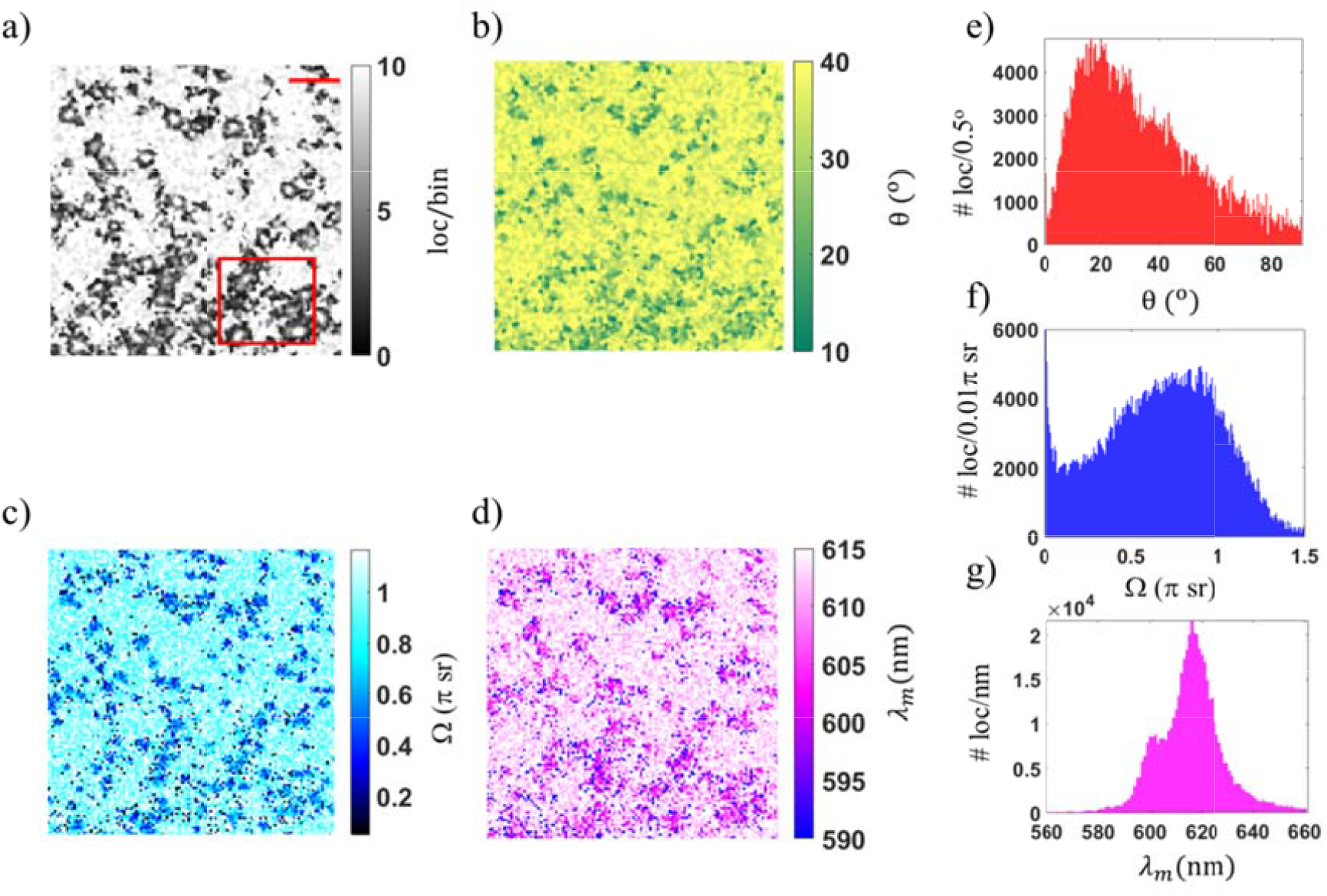
SR-SMOLM images for binding-activated PAINT of NR on DOPC/DPPC/Cholesterol SLB for the same region, where (a) is SMLM image, (b) is polar angle (*θ*), (c) is wobbling area (*Ω*) and (d) is spectral centroid (**λ**_***m***_) map. Color codes are according to the mean orientation of dipoles in each bin. Th polar angle map shows nearly vertically oriented dipoles (*θ* close to 0°) with low wobbling area, identical to Lo phase. NR emission at lower wavelengths from the same region, expected from less hydrated region coinciding with predominant cholesterol so ubilization region, further corroborates above inference. On the contrary, molecules showing red shifted emission with relatively less reorientation constraint and larger polar angle indicates Ld phase of SLB. The azimuthal (*ϕ*) angle map is not relevant in the present context and hence not shown. Distribution of (e) polar angle, (f) wobbling area and (g) spectral centroid observed from their corresponding images. Two distinct sub-population in the spectral centroid distribution in addition to non-uniform orientation parasmeter maps suggest nanoscopic segregation of lipid domains. Scale bar: 1 *μ*m.

NR is reported to occupy the interfacial region of the membrane similar to cholesterol. The influence of cholesterol on the lipid organization within the bilayer, along with the non-covalent interactions between the flat 4-ring structure of cholesterol and the planar benzophenoxazine moiety of NR, guides the later to align itself in accordance with the orientation of cholesterol. Recently, Lew & coworkers^20, 21, 31^ have reported nearly vertical dipole orientation for NR molecules in SLB. This alignment effect is notably strong, as evidenced by a significant reduction in the polar angle (*cf*. **Figure 2b, 3b**) of NR in the Lo phase due to rich presence of cholesterol and DPPC. In the Lo phase, NR exhibited a polar angle distribution around 15°, closely resembling the vertically oriented dipole arrangement seen in literature. In contrast, the Ld phase with dominant presence of DOPC, with two unsaturated acyl chains, exhibited a polar angle distribution close to 30°. For the same reason, NR being solvatochromic in nature, shows blue shift in spectral centroid in Lo phase as compared to Ld phase (*cf*. **Figure 2d, 3d**), further corroborates the dominant presence of cholesterol in the Lo phase, which is in accordance with previous studies.

Earlier works suggest that the orientation map of NR, including polar angle and wobble, are more strongly influenced by specific lipid acyl chains rather than headgroups. Due to the tighter packing in the Lo phase compared to the Ld phase, the wobbling area map (*cf*. **Figures 2c, 3c**) showed a distribution around 0.9 π sr in Lo compared to 1.4 π sr in Ld. These observed trends of polar angle and wobbling area obtained using Vortexed PSF matches well with earlier SMOLM reports with polarized SM PSFs.^20, 21, 31^

Further examination of results (i.e. **Figure 3a-d**) following principal component analysis (PCA) reveals visible separation of Lo and Ld phases (**Figure 3e**, boundary marked by green or black lines). Even a closer look at all the physicochemical parameters to describe NR on membrane along the marked diagonal (shown in **Figure 3a-e**) consistently epitomizes the correlation between lipid order and polarity (*cf*. **Figure 3f**). Parameter distribution of *θ, Ω* and *λ*_m_ for the delineated Lo and Ld regions in **Figure 3a,e**, shown in **Figure 3g**, clearly indicate differential molecular interaction for NR in Lo and Ld phases, as expected and reported in literature. However, in order to establish correlation between polarity and order, we plotted *θ* and *Ω* as a function of *λ*_m_, shown in **Figure 3h**. Striking correspondence between wobbling area and spectral centroid (*cf*. **Figure 3h(ii)**), namely higher rotational mobility (*Ω>*1 *π* sr) due to disorder structure in the cholesterol poor region (Ld) results higher polarity (*λ*_m_∼620 nm) owing to greater hydration by water molecules and the relatively lower mobility (*Ω≤*0.5 *π* sr) due to structural order in the cholesterol rich region (Lo) effects lower polarity (*λ*_m_∼600 nm) by expulsion of water molecules demonstrate the power of SR-SMOLM. Similar connection between polar angle and spectral centroid (*cf*. **Figure 3h(i)**) though present but not very pronounced. For a visual guide to illustrate SR-SMOLM, SM Vortexed PSF with its dispersed spectral PSF for two representative NR molecule from Lo (*θ*=10°, *Ω*=0.3 *π* sr, *λ*_m_=602 nm) and Ld (*θ*=35°, *Ω*=0.8 *π* sr, *λ*_m_=617 nm) region along with their constructed emission spectra are shown in **Figure 3i,j**.

**Figure 3.**
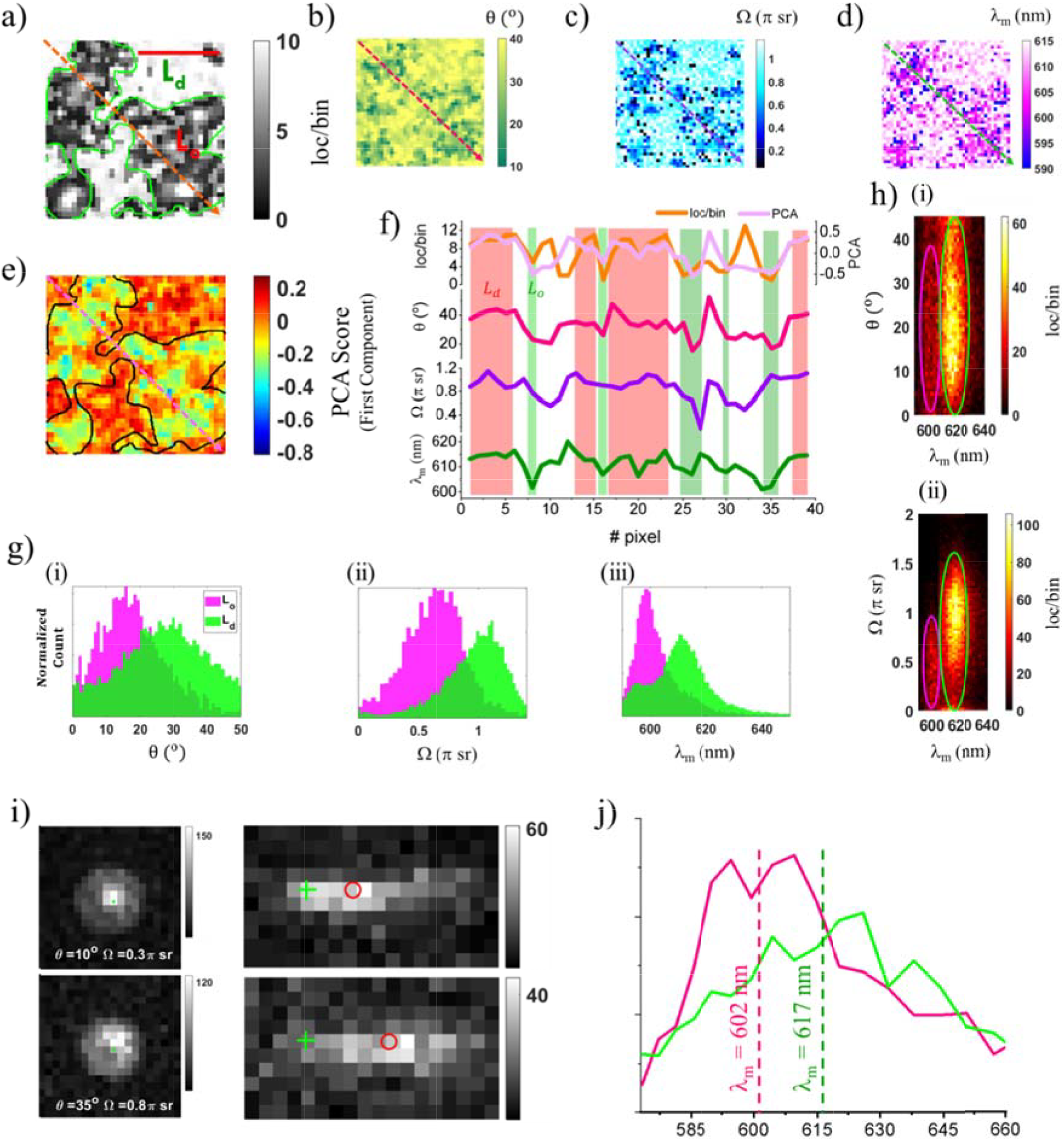
Localization (a) orientation (θ,Ω) and Spectral centroid (*λ*_m_) map of a small area marked with a red square in Figure 2 is further analyzed with PCA (e,f). Boundaries are marked for better visualization of Lo and Ld regions in (a,e). All the parameters along the red dotted line illustrate connection among the extracted values (f). Separate distribution plots (g) for θ, Ω and *λ*_m_ highlight distinct local environment for Lo and Ld regions. Correlation between polarity (*λ*_m_) and order (*θ* and *Ω*) can be easily concluded (h). Representative Vortexed PSF and spectral dispersion (i) of two SMs for Lo and Ld regions along with their calculated spectra (j) clearly highlight effectiveness of SR-SMOLM in addressing micro- heterogeneities. Scale bar: 1 *μ*m.

In conclusion, employing SR-SMOLM we could directly demonstrate the correlation between membrane polarity and order by simultaneous measurement of spectral position and rotational constraint using a single environment sensitive probe with SM sensitivity and high localization precision. The proposed SR-SMOLM is a very powerful tool to explore nanoscale heterogeneity and dynamics of cell membrane with never seen before correlation between structural order and local reaction field which may or may not go hand in hand depending on involved membrane machinery and context.

## ASSOCIATED CONTENT

### Supporting Information

Experimental setup & methods, SM spectral and Vortexed PSF analysis, simulation results for accuracy & precision of extracted parameters, and supplementary figures for NR SRM.

## AUTHOR INFORMATION

### Notes

The authors declare no competing financial interests.

## ACKNOWLEDGMENT

This work is funded and supported by BARC (DAE), India.

