## Supplementary Information for "Spectrally Resolved Single Molecule Orientation Imaging Reveals Direct Correspondence between Polarity and Order Experienced by Nile Red in Supported Lipid Bilayer Membrane"

**1. Materials**:

1,2-dioleoyl-sn-glycero-3-phosphocholine (DOPC) and 1,2-dipalmitoyl-sn-glycero-3-phosphocholine (DPPC) were purchased from Avanti Polar Lipids (Alabaster, AL). Cholesterol was purchased from Sigma Aldrich (St. Louis, Mo). Merocyanine 540 (MC540) and Nile Red were sourced from Thermo Fisher Scientific. HPLC grade water were obtained from Merck Life Science Pvt. Ltd. Solvent DMSO (HPLC grade) was procured from Fisher Scientific. All chemicals were used as received without further purification.

Cell membranes are characterized by their heterogeneous surfaces, consisting of clusters of molecules that form distinct membrane domains. In eukaryotic cells, cholesterol plays a crucial role in driving the formation of these lateral inhomogeneities. To understand these complex domains, simplified model systems with a limited number of lipid types have been used. For the coexistence of liquid-ordered (Lo) and liquid-disordered (Ld) phases, a ternary system is required, with cholesterol being an essential component. The remaining two lipids are selected to have significantly different main phase transition temperatures ($T_m$). Hence, the obvious choice is an unsaturated lipid with a lower $T_m$ and a saturated lipid with a higher $T_m$. These model ternary systems exhibit liquid–liquid phase coexistence, forming two distinct phases: the liquid-ordered phase (Lo), enriched in high-melting lipids and cholesterol and thought to resemble the composition of lipid rafts, and the liquid-disordered phase (Ld), which surrounds the rafts. Consequently, we have selected a ternary mixture comprising the unsaturated lipid DOPC ($T_m$ = -17°C[1]), the saturated lipid DPPC ($T_m$ = 41°C[2]), and cholesterol, a combination known to demonstrate the existence of Lo and Ld phases.

### 2. Imaging buffer

Tris buffer: Comprising 10 mM Tris, 100 mM NaCl, pH 7.4 is used for SR-SMOLM of Nile red. GLOX buffer: Comprising 10 mM NaCl, 10% (w/v) glucose, and 1% (v/v) enzymatic oxygen scavenger system in 50 mM Tris (pH 8.3) is used for SMLM imaging with MC540. The enzymatic oxygen scavenger system stock solution was prepared by combining 8 mg of glucose oxidase and 38 µl of a 21 mg/mL catalase solution with 160 µl of PBS. After centrifugation at 15,000 rpm for 1 minute, the precipitate was discarded prior to use.

Nanomolar dye solutions in water were prepared from a concentrated dye stock (in DMSO-water). Around 50 µl was added on to SLB membrane for SR-SMOLM imaging and suitably adjusted the blinking SM density using blank buffer.

### 3. SLB preparation

**i) Large unilamellar vesicles (LUVs):** The LUVs were prepared by the extrusion method. Briefly, the required amount of DOPC, DPPC and cholesterol were dissolved in chloroform to achieve a molar ratio of 35:35:30. Lipid films were obtained by first using $N_2$ and then vacuum evaporation of organic solvent.[3] These films were hydrated in tris buffer (10 mM tris, 100 mM NaCl, 3 mM $CaCl_2$, pH 7.4) to achieve a concentration of 2 mM for DOPC and DPPC. Lipid suspensions were subjected to freeze−thaw cycles, 10 times (from -80 °C to room temperature) with intermittent mixing, to give multilamellar vesicles (MLVs). MLVs were extruded 20 times through a polycarbonate filter of 100 nm pore size using a mini extruder from Avanti Polar Lipids Inc at room temperature.

**ii) Supported lipid bilayer (SLB):** To prepare SLBs, LUVs were added onto clean coverslips and incubated at 55°C for 1 hour to form a SLB. Subsequently, unfused vesicles were removed by washing with HPLC grade water (at 50 °C) by carefully pipetting in and out. After 30 min of cooling to room temperature, the lipid bilayer was thoroughly rinsed with Tris buffer to remove any residual lipids and imaged immediately.

**iii) Cover slip cleaning:** Glass cover slides (thickness 1.5) were first cleaned with 1x Hellmanex detergent solution and then rinsed with de-ionized water several times. The rinsed slides were the sonicated with 2M $H_2SO_4$ for 30 mins, rinsed few times with de-ionized water and then further sonication in de-ionized water for 30 mins. The slides were then rinsed with HPLC grade ethanol and dried. Slides were used within seven days of cleaning.

### 4. SMLM and SRSMOLM imaging in SLB

The lipophilic dyes, such as MC540 and Nile Red temporarily adhere to lipid bilayers and are suitable for PAINT-based SMLM and SMOLM techniques. These dyes are non-fluorescent in aqueous media; radiative de-excitation gets activated upon binding to membrane. The blinking frequency of these probes was regulated by adjusting their concentrations in the imaging buffer and/or the intensity of the incident laser to ensure the observation of single-molecule fluorescence without substantial overlap of PSFs. Initially, MC540 imaging was conducted. A sequence of 45,000 images was recorded with 50 ms exposure time and 2 kW/cm$^2$ excitation power density.

Subsequently, the SLB sample underwent careful and thorough rinsing with Tris buffer prior to addition of Nile Red for imaging.

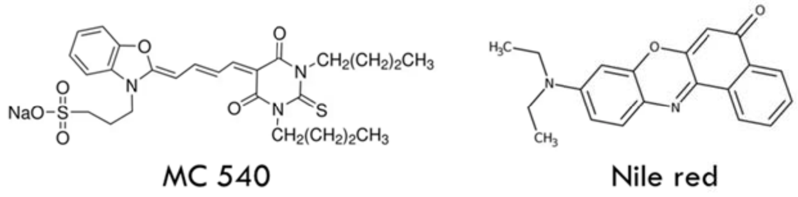


**Scheme S1.** Chemical structure of membrane probes Nile Red and MC 540.

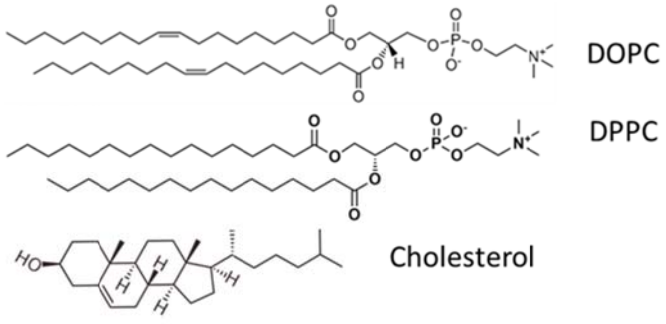


**Scheme S2.** Chemical structure of lipids used to in SLB preparation.

### 5. SR-SMOLM setup

The single molecule (SM) imaging experiments were conducted utilizing a custom-built wide-field epifluorescence microscope. In the excitation phase, a circularly polarized excitation beam with an intensity of approximately 2 kW/cm² was directed onto the back focal plane of the objective lens (100x NA1.46 oil immersion, Carl Zeiss GmbH) to ensure uniform total internal reflection (TIR) illumination of the sample. Fluorescence emission was captured through a combination of a dichroic mirror (FF538-FDi01, Semrock) and filters (532 Notch filter, Semrock, 561 nm Long Pass, Semrock, and a 650 nm Short Pass, Semrock).The detection system comprised two distinct paths, labeled as Path 1 and Path 2, which were separated using a 70:30 beam splitter (Thorlabs BS081 - 30:70 (R:T)). Path 1 was dedicated to localization-orientation microscopy employing a vortex waveplate (V-593-10-1, vortex photonics) positioned at the Fourier plane (FP) of the 4*f*-imaging system,[4] resulting in a magnification factor of 2x and achieving a pixel size of 80 nm. Conversely, Path 2 facilitated spectral imaging through the utilization of a dispersive glass prism also placed at the Fourier plane. For this path no additional magnification is employed to resists further photon loss. Both emission channels were simultaneously recorded using a single Newton EMCCD camera (DU971N-UVB, Andor). Image acquisition involved recording image stacks with a 50 ms exposure time, comprising 45,000 sequential frames. Orientation-localization microscopy (SMOLM), based on the principle of points accumulation for imaging in nanoscale topography (PAINT), achieved super-resolution by isolating individual emitters through fitting the point spread function (PSF) using either Gauss MLE or weighted least-square estimator. The former estimator offered superior precision compared to the latter. The concentration of probes utilized in SMLM experiments ranged from 10 to 80 nM. Subsequently, recorded images underwent post-processing for drift correction and density filtering using custom MATLAB scripts.

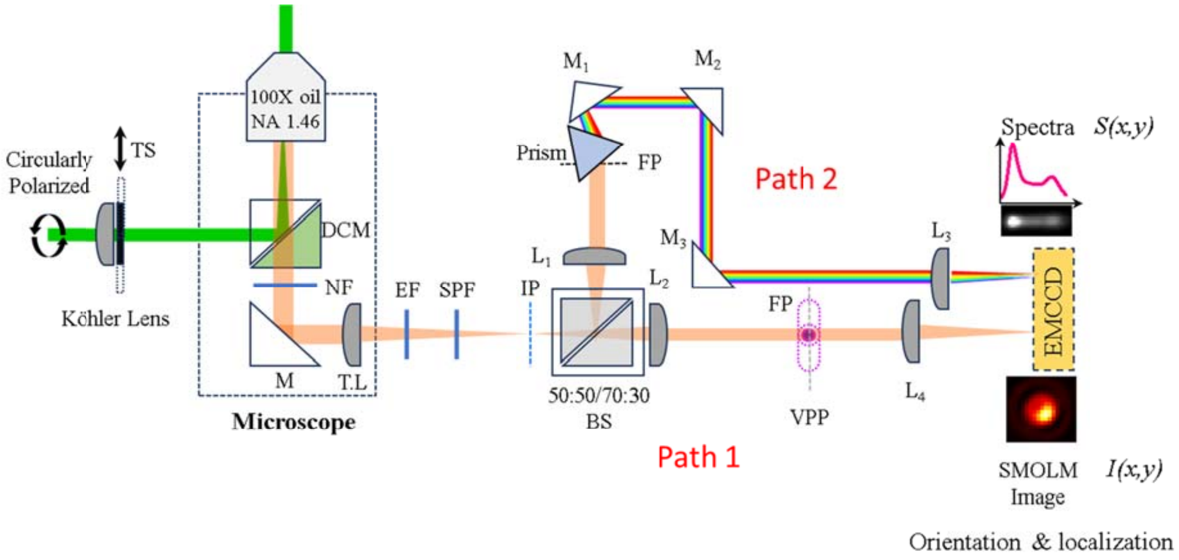


**Figure S1:** Schematic diagram of SR-SMOLM setup. Path 1 is used for SMOLM using Vortex phase plate (VPP) placed at BFP of the 4*f* system. Path 2 is used for simultaneous spectral imaging using a prism placed at Fourier plane (FP). TS: Translational stage, M: mirrors, DCM: Dichroic mirror, NF: Notch Filter, TL: Tube lens, EF: Emission Filter, SPF: Short pass filter, IP: Image plane, L: Lenses, FP: Fourier plane, BS: Beam splitter, VPP: Vortex phase-plate.

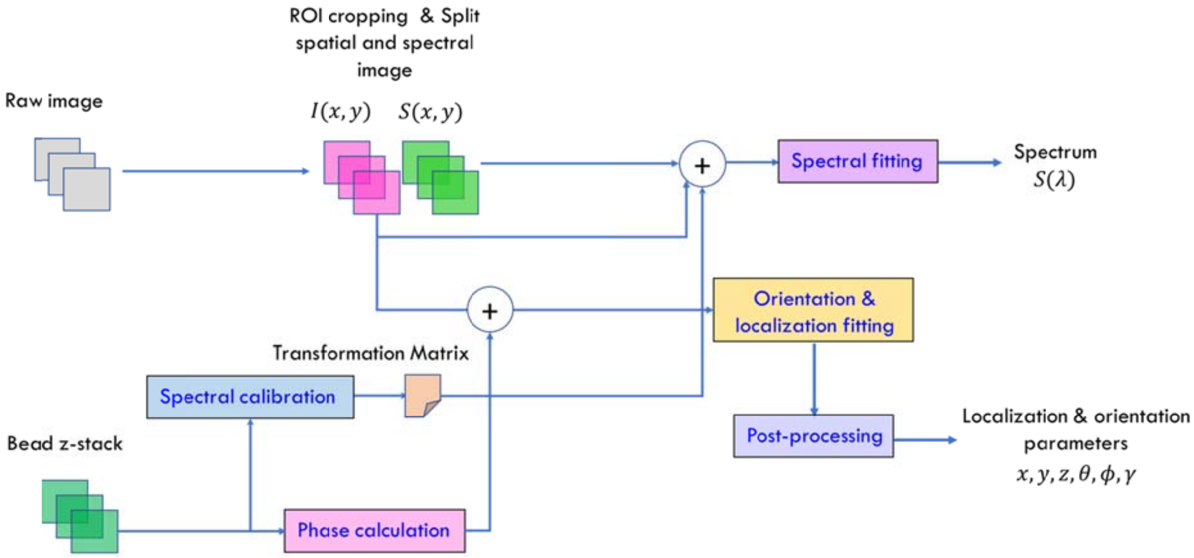


**Schematic S3**: Work flow diagram of the image processing for SR-SMOLM.

The initial phase entails partitioning the captured raw image stack into distinct Regions of Interest (ROIs) for both spatial (I(x,y)) and spectral (S(x,y)) images. The spatial image serves the purpose of extracting localization and orientation parameters, whereas the spectral image is instrumental in computing the spectra of identified molecules within the spatial domain.

To derive localization and orientation parameters, we have adopted the methodology elucidated by Hulleman et al.[4] A tailored Point Spread Function (PSF) model is formulated, encompassing a zone function that corresponds to the Vortex Phase Plate (VPP), alongside additional aberrations arising from misalignment and imperfections within the optical apparatus. The calculation of aberrations employs a Zernike polynomial-based methodology, utilizing z-stack images of fluorescent beads (Tetraspeck, 100 nm) for calibration purposes.

The spectral image[5-10] undergoes processing utilizing two pre-generated calibration files: The first file facilitates the creation of a spatial-to-spectral mapping (achieved through a polynomial transformation of degree-2 using fitgeotrans in MATLAB), correlating the spatial image with the corresponding spectral position of 590±10 nm emission via a narrow bandpass. The second calibration file is generated through a sequence of similar narrow bandpass filters to establish the calibration curve between pixel shift and the corresponding wavelength value.

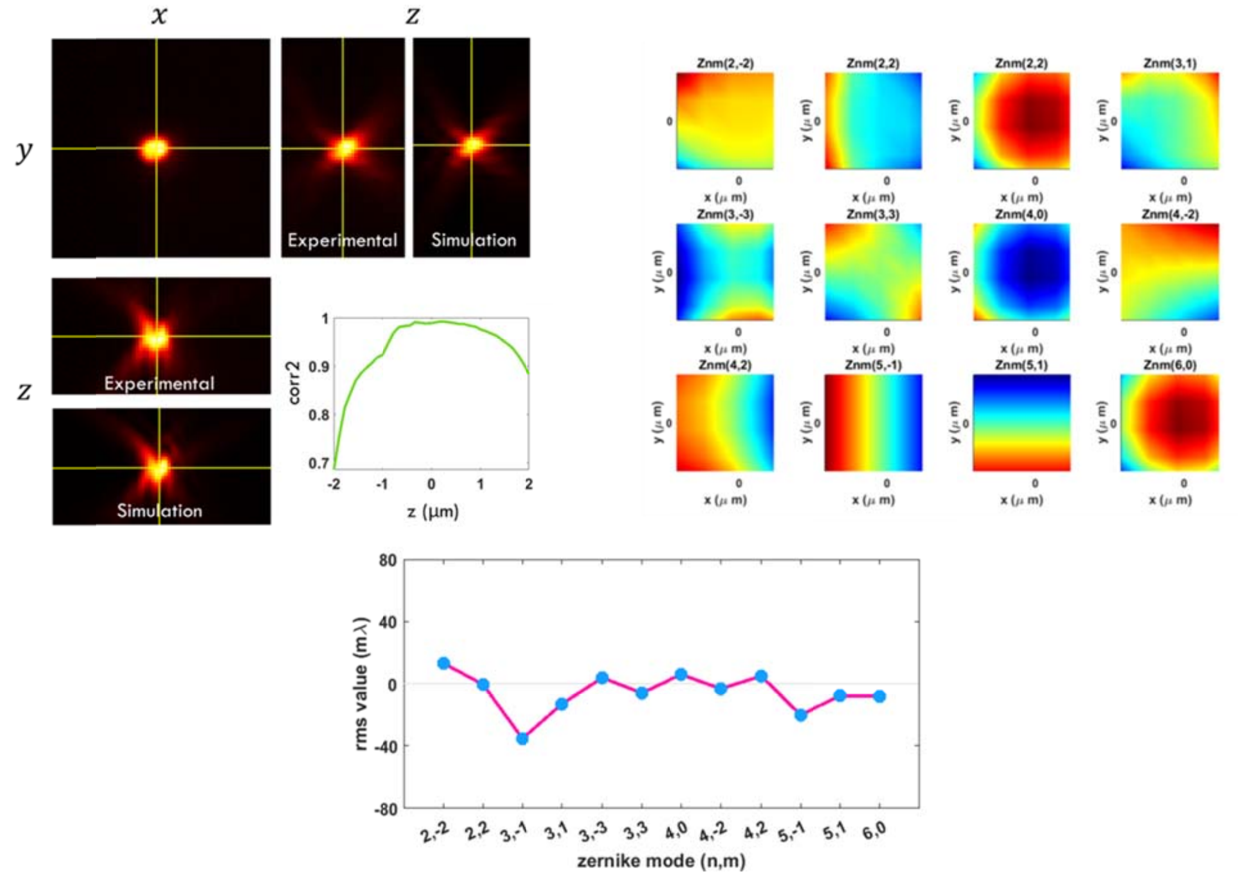


**Figure S2:** On the top left – experimental and simulated z-stack profile of a Tetraspeck bead in xz- and yz-plane with corresponding correlation. On the right- calculated 2D surface plots of Zernike coefficients (Blue-valley, Red -Peak). On the bottom – rms value of different Zernike coefficients

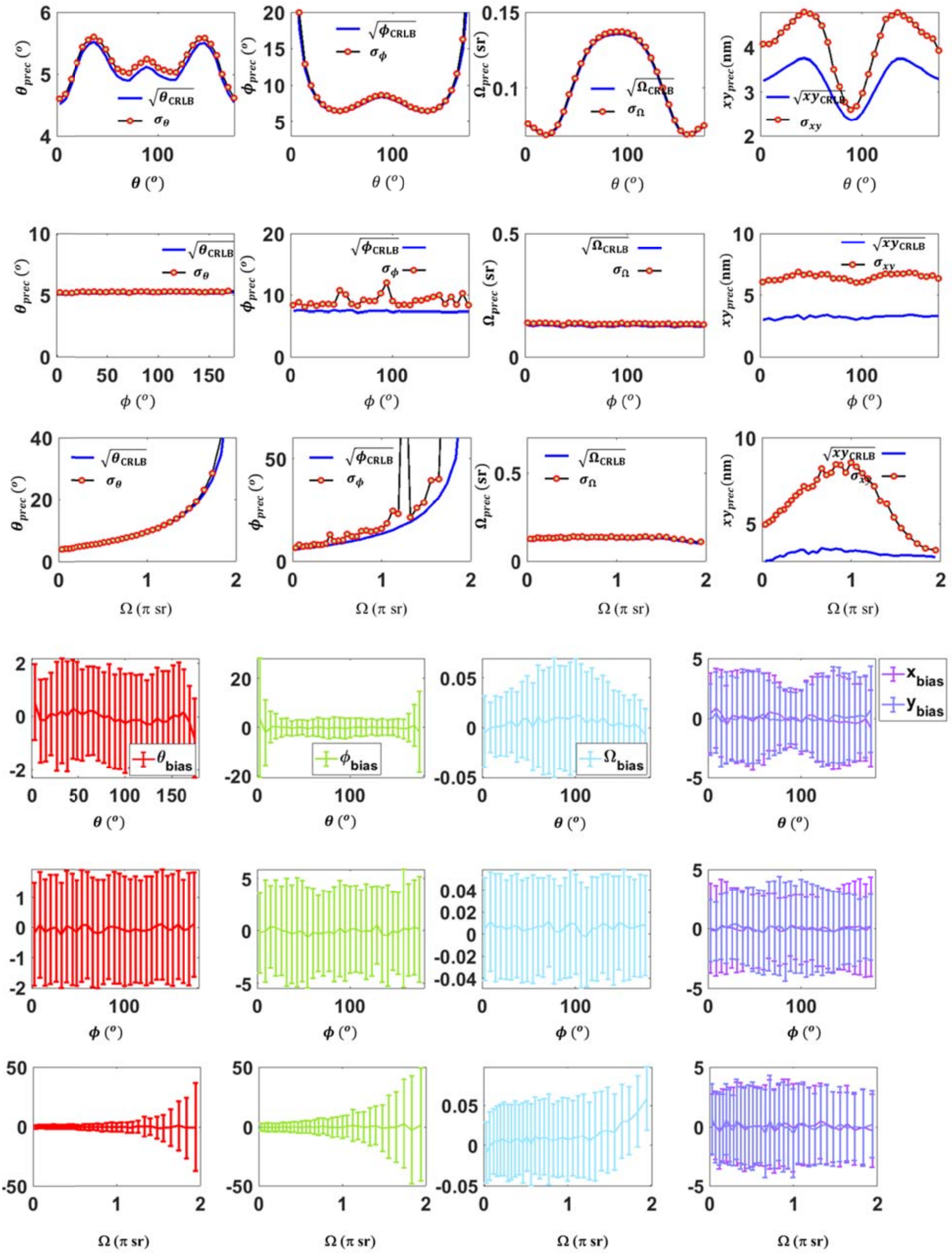

**Figure S3:** Simulation results showing the performance of our PSF fitting in terms of precision and the bias of different associated orientation-localization parameters.

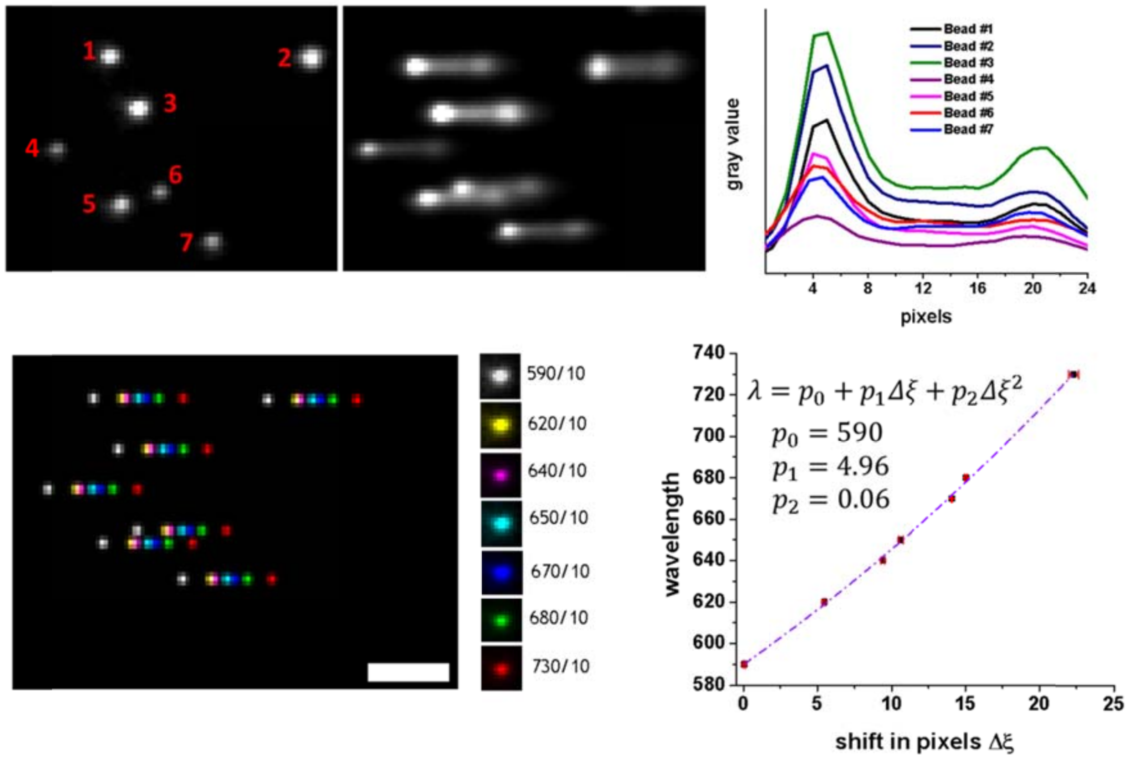


**Figure S4:** Top panel – SMOLM images of Tetraspeck beads (on the left, numbered) with corresponding spectrally dispersed PSFs (middle) and individual spectra (right). Bottom panel – composite image shows spectra profile after employing 7 different narrow bandpass filters (Thorlabs, FBH590-10, FBH620-10, FBH640-10, FBH650-10, FBH650-10, FBH670-10, FBH680-10 & FBH730-10) (on the left) and corresponding wavelength – pixel-shift characteristic curve fitted with a second order polynomial. Scale bar: 1 μm.
